## Supplementary Information for "Temperature-dependence of Early Development of Zebrafish and the Consequences for Laboratory Use and Animal Welfare"

#### 8 Supplementary Information

##### Table of Contents

*8.1 Embryonic Stages Categorization* 2

*8.2 Developmental Delay of All Endpoints by Experimental Replicate* 3

*8.3 Statistical parameters and diagnostic plots for the High-Resolution and Low-Resolution Time Series* 4

*8.4 Delay of Early Development Stages* 7

*8.5 Comparison of Current Findings with Kimmel et al. (1995)* 9

*8.6 Reanalysis of Kimmel et al. (1995) Data* 10

#### 8.1 Embryonic Stages Categorization

***Table S1**: Descriptions of the embryonic stages determined during the high-resolution time series. Stages were categorized according to the developmental criteria of Kimmel et al. (1995). The table shows the selection of criteria from Kimmel et al. (1995 ) used to determine the respective stage.*

| Stage | Abbreviation | Description |
|---|---|---|
| 4 cell | 4c | 2 x 2 array of blastomeres |
| 8 cell | 8c | 2 x 4 array of blastomeres |
| 16 cell | 16c | 4 x 4 array of blastomeres |
| 32 cell | 32c | 2 regular tier (horizontal rows) of blastomeres, 4 x 8 array |
| 64 cell | 64 | 3 regular tier of blastomeres |
| 128 cell | 128 | 5 blastomeres tier, cleavage planes irregular |
| 256 cell | 256c | 7 - 9 irregular blastomeres tiers, yolk syncytial layer develops |
| oblong | O | > = 11 irregular blastomeres tiers, beginning of blastodisc cell asynchrony and flattening. Flattening produces an elliptical shape. Animal-vegetal axis of the blastula shortens, with the blastodisc compressing down upon the yolk cell |
| sphere | S | Continued shortening along tha animla-vegetal axis generates a late blastula of smooth and approximately spherical shape, flat border between blastodisc and yolk, start of yolk cell bulging (doming) toward animal pole as epiboly begins |
| epiboly | E | Epiboly is the thinning and spreading of the yolk syncytical layer and the blastodisc over the yolk cell. 30%-50% epiboly. |
| shield | SH | germ ring/ shield visible from animal pole. After reaching a 50% epiboly a thickened marginal region (germ ring), nearly simultaneously all around the blastoderm rim. Convergence movements then, nearly as rapidly, produce a local accumulation of cells at one position along the germ ring, the embryonic shield |
| 75 epiboly | 75E | Dorsal side distinctly thicker, thin evacuation zone on ventral side |
| 90 epiboly | 90E | Brain rudiment thickened, notochord rudiment distinct from segmental plate |
| bud | B | tail bud prominent, early polster, along dorsal side the neural plate is thickened along the embryonic axis. The thickening is most prominent near the animal pole in the prospective head region, where the head will form |
| 3 somite | 3s | first three somites furrow |
| 6 somite | 6s | brain primordium has distinctively thickened, eye primordium |
| 8 somite | 8s | 8 - 10 somites without otic placode |
| 10 somite | 10s | otic placode developed |
| 14 somite | 14s | yolk cell begins to invert and look like kidney-bean. Somites begin to take on chevron shape |
| 18 somite | 18s | yolk cell extension is clearly delimited from the yolk ball as the tail straightens out |
| 21 somite | 21s | lens primordium |
| 26 somite | 26s | Straightening of posterior trunk is nearly completed, but elongating tail still curves ventrally. Cerebellar primordium is prominent |
| prim | P | Cerebellum is evident at the hindbrain/midbrain boundary region |
| Eye pigmentation | EP | eye pigmentation begins |
| Body pigmentation | BP | body pigmentation begins |

#### *8.2 Developmental Delay of All Endpoints by Experimental Replicate*

***Table S2:*** *Delays in development observed for different endpoints computed with confidence interval separately for each replicate. The coefficient of variation between replicates is given for all endpoints.*

| | | **Hours of delay** | | |
|---|---|---|---|---|
| **Endpoint** | dpf | min | average | max |
| **Early Developmental Stages (Body Pigmentation (BP)*)** | 1 | | | |
| R1 | | 3 | 4 | 4.5 |
| R2 | | 1 | 3 | 5 |
| Coefficient of Variation | | 0.71 | 0.20 | 0.07 |
| **Onset of Heartbeat** | 2 | | | |
| R1 | | 3 | 3 | 5 |
| R2 | | 4 | 5.5 | 6 |
| Coefficient of Variation | | 0.20 | 0.42 | 0.13 |
| **Hatching** | 3 | | | |
| R1 | | 3.5 | 5 | 11.5 |
| R2 | | 1 | 6 | 14 |
| Coefficient of Variation | | 0.79 | 0.13 | 0.14 |
| **Body Length hr** | 5 | | | |
| R1 | | -6 | 3.4 | 10 |
| R2 | | 17.8 | 22 | 25.6 |
| Coefficient of Variation | | 2.85 | 1.04 | 0.62 |
| **Body Length lr** | 5 | | | |
| R1 | | 4.6 | 12.9 | 19.1 |
| R2 | | 5 | 10.7 | 15.6 |
| R3 | | -0.8 | 12.3 | 18.8 |
| Coefficient of Variation | | 1.10 | 0.10 | 0.11 |
| **Eye Size** | 5 | | | |
| R1 | | -1.5 | 8.26 | 19.5 |
| R2 | | 7.1 | 17.9 | 28.1 |
| R3 | | 6.6 | 16.7 | 27.4 |
| Coefficient of Variation | | 1.19 | 0.37 | 0.19 |
| **Yolk Sac Consumption** | 5 | | | |
| R1 | | 12.4 | 22.4 | 31.3 |
| R2 | | 6.8 | 13.2 | 18.5 |
| R3 | | 7 | 16.8 | 25.6 |
| Coefficient of Variation | | 0.36 | 0.27 | 0.26 |

*the final stage observed in the early developmental period and its associated temporal endpoint.
hr = high-resolution time series, lr = low-resolution time series

*8.3 Statistical parameters and diagnostic plots for the High-Resolution and Low-Resolution Time Series*

### Body length

#### High-resolution 26 °C

| | Estimate | Std. Error | t value | Pr(>\|t\|) |
|---|---|---|---|---|
| (Intercept) | 2.95774 | 0.03321 | 89.06 | <2e-16 *** |
| Approximate significance of smooth terms: | | | | |
| | edf | Ref.df | F | p-value |
| s(hpf) | 3 | 3.001 | 1486.5 | <2e-16 *** |
| s(replicate) | 0.992 | 1 | 358.7 | <2e-16 *** |
| R-sq.(adj) = 0.7935 | Deviance explained = 79.4% | | | |
| GCV score = 0.011127 | Scale est. = 0.011083 | n = 1262 | | |

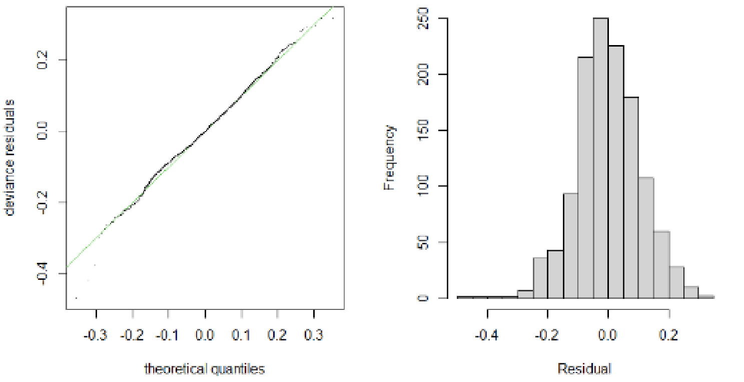


#### High-resolution 28 °C

| | Estimate | Std. Error | t value | Pr(>\|t\|) |
|---|---|---|---|---|
| (Intercept) | 3.03041 | 0.02754 | 110 | <2e-16 *** |
| Approximate significance of smooth terms: | | | | |
| | edf | Ref.df | F | p-value |
| s(hpf) | 3.0294 | 3.056 | 3463.69 | <2e-16 *** |
| s(replicate) | 0.9935 | 1 | 44.71 | 3.06e-11 *** |
| R-sq.(adj) = 0.8644 | Deviance explained = 86.5% | | | |
| GCV score = 0.0082589 | Scale est. = 0.0082344 | n = 1692 | | |

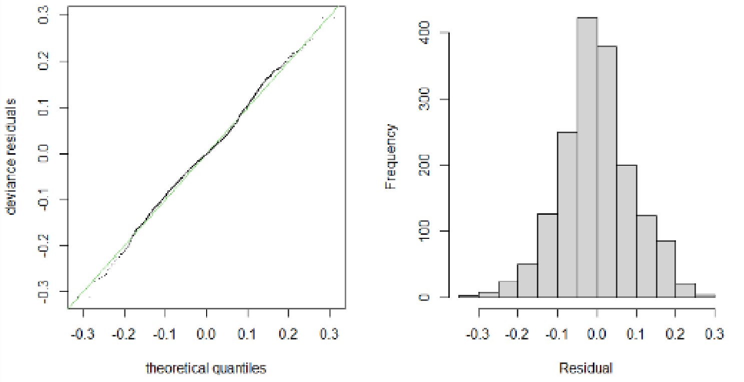


#### Low-resolution 26 °C

| | Estimate | Std. Error | t value | Pr(>\|t\|) |
|---|---|---|---|---|
| (Intercept) | 3.54389 | 0.01541 | 229.9 | <2e-16 *** |
| Approximate significance of smooth terms: | | | | |
| | edf | Ref.df | F | p-value |
| s(hpf) | 2 | 2 | 477.57 | <2e-16 *** |
| s(replicate) | 1.561 | 2 | 5.102 | 0.00551 ** |
| R-sq.(adj) = 0.8491 | Deviance explained = 85.2% | | | |
| GCV score = 0.0090238 | Scale est. = 0.0087845 | n = 172 | | |

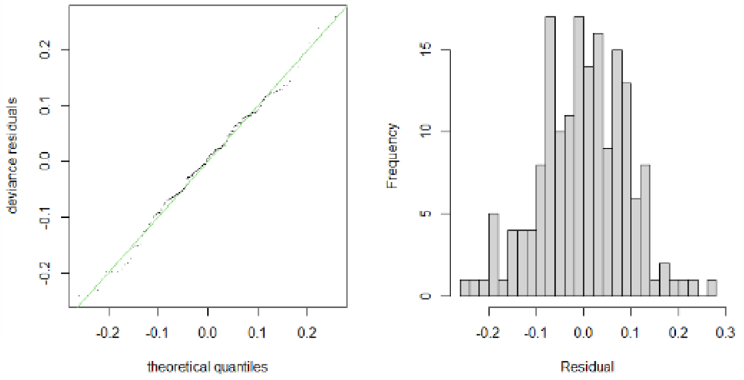


#### Low-resolution 28 °C

| | Estimate | Std. Error | t value | Pr(>\|t\|) |
|---|---|---|---|---|
| (Intercept) | 3.65587 | 0.01612 | 226.8 | <2e-16 *** |
| Approximate significance of smooth terms: | | | | |
| | edf | Ref.df | F | p-value |
| s(hpf) | 2 | 2 | 505.038 | <2e-16 *** |
| s(replicate) | 1.624 | 2 | 5.158 | 0.0056 ** |
| R-sq.(adj) = 0.8577 | Deviance explained = 86.1% | | | |
| GCV score = 0.008534 | Scale est. = 0.0083019 | n = 170 | | |

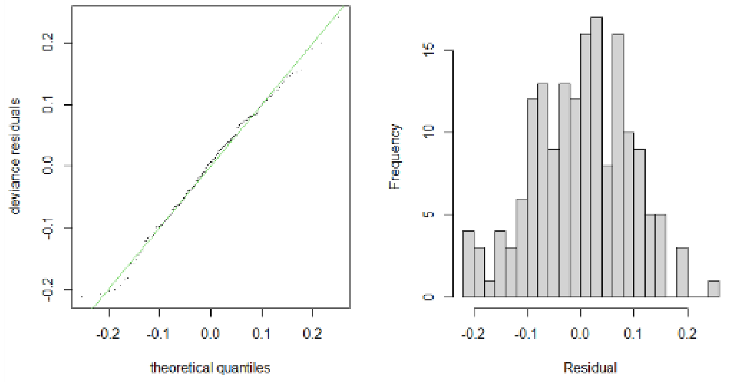


***Fig. S1:*** *Parameters and diagnostic plots for non-parametric Shape Constrained Additive Model (SCAM with integrated smoothness) with a random effects term for replicate.*

| | Estimate | Std.Error | t value | Pr(>\|t\|) |
|---|---|---|---|---|
| (Intercept) | -15.7718 | 1.358748 | -11.61 | <2e-16 *** |
| temperature | 1.404162 | 0.101353 | 13.85 | <2e-16 *** |
| I(temperature^2) | -0.02507 | 0.001878 | -13.35 | <2e-16 *** |
| Residual standard error: 0.1206 on 256 degrees of freedom | | | | |
| Multiple R-squared: 0.6277, Adjusted R-squared: 0.6248 |  |  |  |  |
| F-statistic: 215.8 on 2 and 256 DF, p-value: < 2.2e-16 |  |  |  |  |

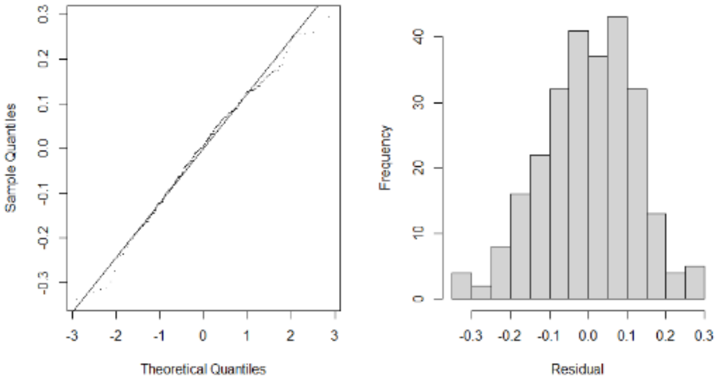


***Fig. S2:*** *Model diagnostics for the regression of body length over temperature with a quadratic model.*

### Yolk sac

#### Low-resolution 26 °C

| Parametric coefficients: | | | | |
|---|---|---|---|---|
| | Estimate | Std. Error | t value | Pr(>\|t\|) |
| (Intercept) | 0.282512 | 0.004489 | 62.94 | <2e-16 *** |
| Approximate significance of smooth terms: | | | | |
| | edf | Ref.df | F | p-value |
| s(hpf) | 2 | 2 | 119.419 | < 2e-16 *** |
| s(replicate | 1.674 | 2 | 9.375 | 0.000107 *** |
| R-sq.(adj) = 0.5914 Deviance explained = 60% | | | | |
| GCV score = 0.00059032 Scale est. = 0.00057464 n = 176 | | | | |

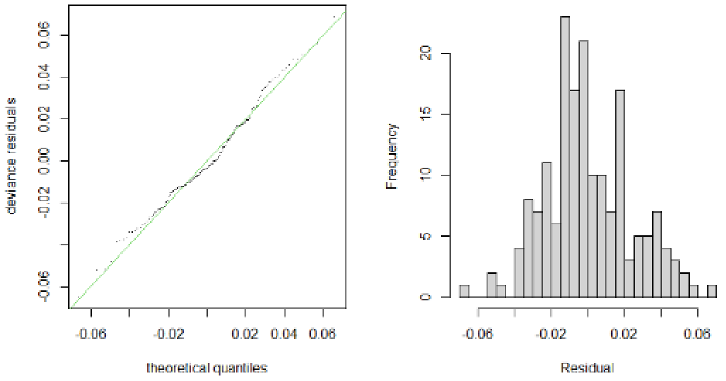


#### Low-resolution 28 °C

| Parametric coefficients: | | | | |
|---|---|---|---|---|
| | Estimate | Std. Error | t value | Pr(>\|t\|) |
| (Intercept) | 0.261565 | 0.002007 | 130.3 | <2e-16 *** |
| Approximate significance of smooth terms: | | | | |
| | edf | Ref.df | F | p-value |
| s(hpf) | 2 | 2 | 155.358 | < 2e-16 *** |
| s(replicate | 0.0133 | 2 | 0.006 | 0.649 |
| R-sq.(adj) = 0.6409 Deviance explained = 64.5% | | | | |
| GCV score = 0.00070851 Scale est. = 0.00069624 n = 174 | | | | |

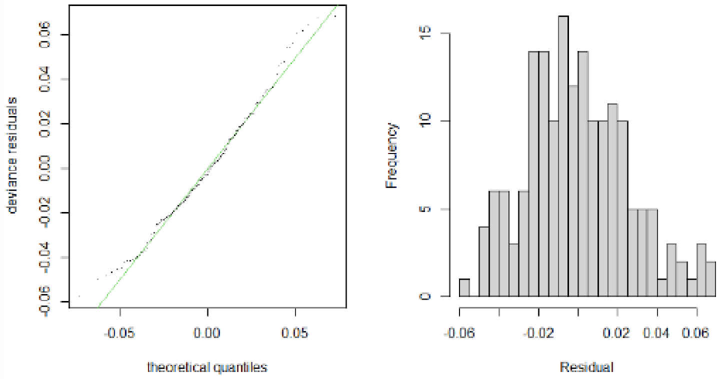


### Eye size

#### Low-resolution 26 °C

| | Estimate | Std. Error | t value | Pr(>\|t\|) |
|---|---|---|---|---|
| (Intercept) | 0.06399 | 0.00252 | 25.39 | <2e-16 *** |
| Approximate significance of smooth terms: | | | | |
| | edf | Ref.df | F | p-value |
| s(hpf) | 2 | 2 | 140.98 | < 2e-16 *** |
| s(replicate) | 1.922 | 2 | 20.14 | 1.16e-08 *** |
| R-sq.(adj) = 0.6413 Deviance explained = 64.9% | | | | |
| GCV score = 4.4e-05 Scale est. = 4.2769e-05 n = 176 | | | | |

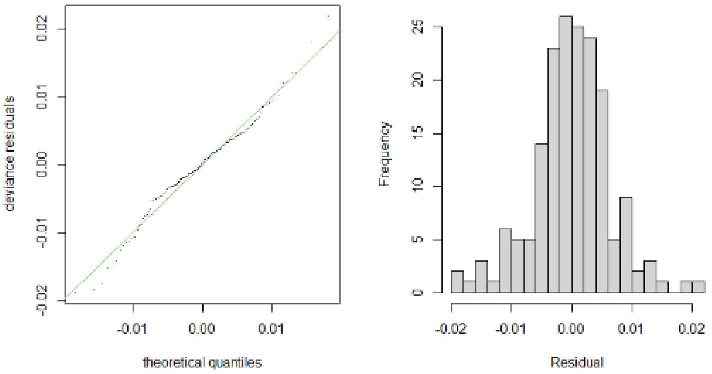


#### Low-resolution 28 °C

| | Estimate | Std. Error | t value | Pr(>\|t\|) |
|---|---|---|---|---|
| (Intercept) | 0.069391 | 0.001438 | 48.27 | <2e-16 *** |
| Approximate significance of smooth terms: | | | | |
| | edf | Ref.df | F | p-value |
| s(hpf) | 2 | 2 | 129.82 | < 2e-16 *** |
| s(replicate) | 1.683 | 2 | 11.21 | 2.02e-05 *** |
| R-sq.(adj) = 0.6166 Deviance explained = 62.5% | | | | |
| GCV score = 5.8763e-05 Scale est. = 5.719e-05 n = 175 | | | | |

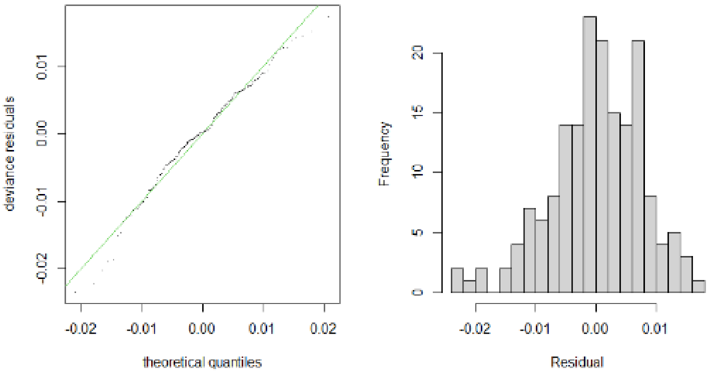


***Fig. S3:*** *Parameters and diagnostic plots for non-parametric Shape Constrained Additive Model (SCAM with integrated smoothness) with a random effects term for replicate*

#### 8.4 Delay of Early Development Stages

***Table S3**: Representative overview of temperature-dependent developmental differences in zebrafish embryos at 26 °C and 28 °C at three early stages: 14, 18, and 24 hpf. The table highlights morphological differences observed at each time point, illustrating the influence of incubation temperature on embryonic development.*

| | **14 hpf** | **18 hpf** | **24 hpf** |
|---|---|---|---|
| **26 °C** | 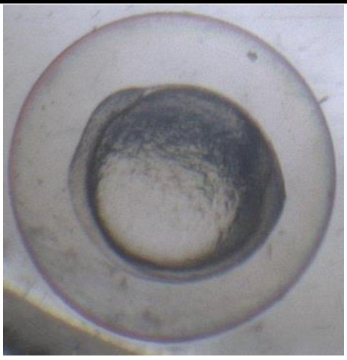 | 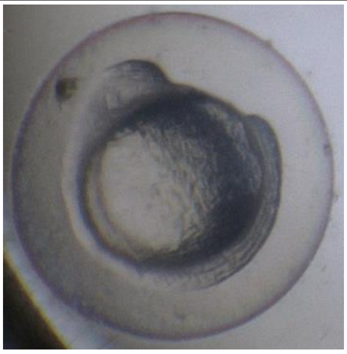 | 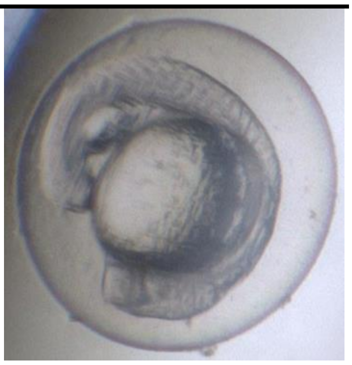 |
| | *Bud* | *8 somite* | *21 somite* |
| **28 °C** | 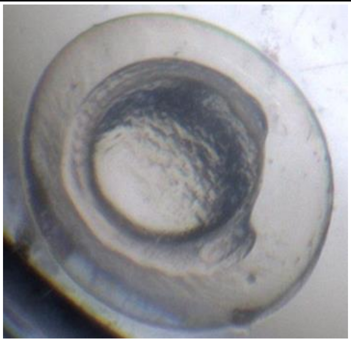 | 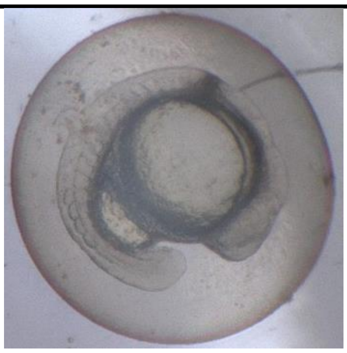 | 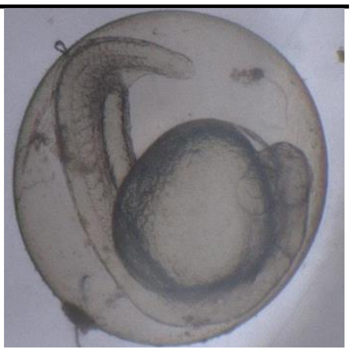 |
| | *8 somite* | *18 somite* | *prim* |

***Table S4**: Delay in hours post fertilization for zebrafish embryos to reach developmental stages at 26 °C and 28 °C. n = number of embryos this stage was observed in, CI = bootstrapped confidence interval of difference in medians confined by replicate, p-value: derived from a permutation test with replicate as blocking factor (2000 resamples) and corrected for alpha inflation with the Bonferroni Method.*

| Stage | Abbreviation | n (26 °C) | n (28 °C) | Difference in medians / h | CI | p-value |
|---|---|---|---|---|---|---|
| 4-cell | 4c | 29 | 20 | 0.0 | 0.0 – 0.0 | - |
| 8-cell | 8c | 20 | 28 | 0.0 | 0.0 – 0.0 | - |
| 16-cell | 16c | 8 | 6 | 0.0 | 0.0 – 0.0 | - |
| 32-cell | 32c | 20 | 34 | 0.0 | 0.0 – 0.0 | - |
| 64-cell | 64c | 14 | 14 | 0.0 | 0.0 – 0.0 | - |
| 128-cell | 128c | 21 | 12 | 0.0 | 0.0 – 0.0 | - |
| 256-cell | 256c | 24 | 33 | 0.5 | 0.0 – 0.0 | - |
| oblong | O | 22 | 26 | 0.0 | 0.0 – 0.0 | - |
| sphere | S | 17 | 18 | 1.0 | 0.0 – 0.0 | - |
| epiboly | E | 28 | 33 | 0.0 | 0.0 – 1.5 | 0.157 |
| shield | SH | 40 | 41 | 1.0 | 0.25 – 1.5 | < 0.0001 |
| 75-epiboly | 75E | 29 | 26 | 1.0 | 1.5 – 2.5 | < 0.0001 |
| 90-epiboly | 90E | 41 | 44 | 2.0 | 0.25 – 2.0 | < 0.0001 |
| bud | B | 43 | 42 | 2.0 | 2.5 – 3.0 | < 0.0001 |
| 3-somite | 3s | 38 | 37 | 2.0 | 2.0 – 3 | < 0.0001 |
| 6-somite | 6s | 34 | 44 | 2.0 | 2.5 – 3.5 | < 0.0001 |
| 8-somite | 8s | 36 | 25 | 3.0 | 2.25 – 3.5 | < 0.0001 |
| 10-somite | 10s | 31 | 31 | 2.0 | 1.5 – 3.5 | < 0.0001 |
| 14-somite | 14s | 24 | 13 | 2.5 | 1.5 – 3.0 | < 0.0001 |
| 18-somite | 18s | 36 | 36 | 3.0 | 2.5 – 4.0 | < 0.0001 |
| 21-somite | 21s | 43 | 41 | 3.0 | 2.75 – 4.0 | < 0.0001 |
| 26-somite | 26s | 37 | 43 | 4.0 | 3.0 – 5.25 | < 0.0001 |
| prim | P | 43 | 39 | 5.0 | 4.25 – 6.25 | < 0.0001 |
| pigmented eyes | EP | 37 | 41 | 4.0 | 4.5 – 6.0 | < 0.0001 |
| pigmented body | BP | 44 | 46 | 3.0 | 3.25 – 5.0 | < 0.0001 |

*8.5 Comparison of Current Findings with Kimmel et al. (1995)*

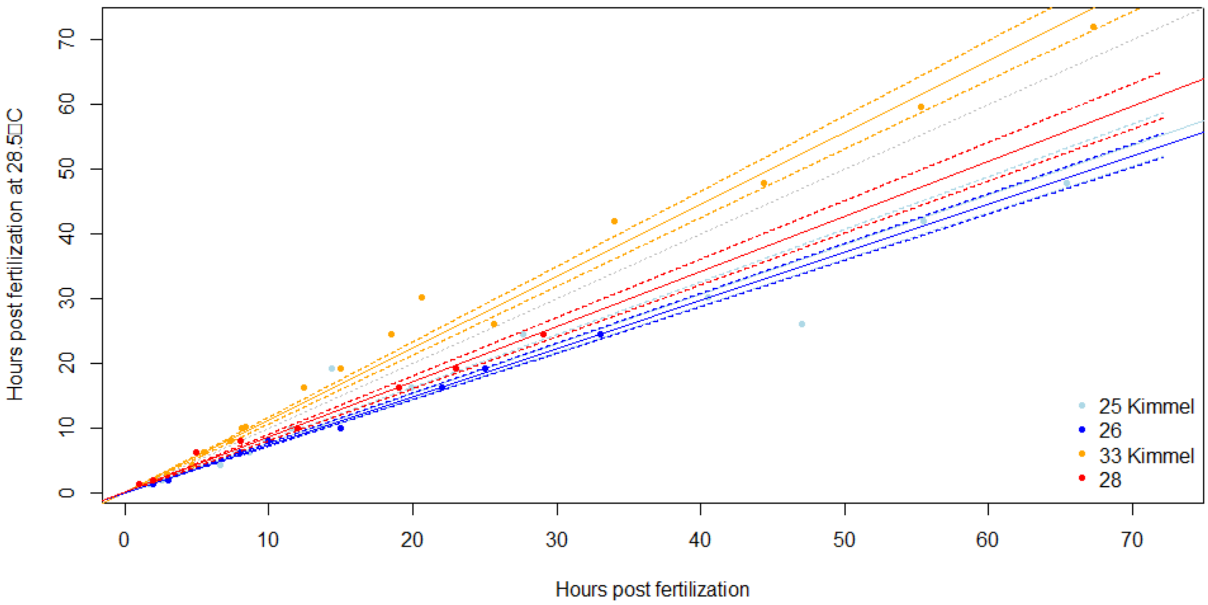


***Fig. S4:*** *Relationship between the hours post fertilization (hpf) at different temperatures and the corresponding normalized hpf at 28.5 °C, comparing data generated on zebrafish embryo stages by Kimmel et al. 1995 to our data on staging, where comparable.*

#### *8.6 Reanalysis of Kimmel et al. (1995) Data*

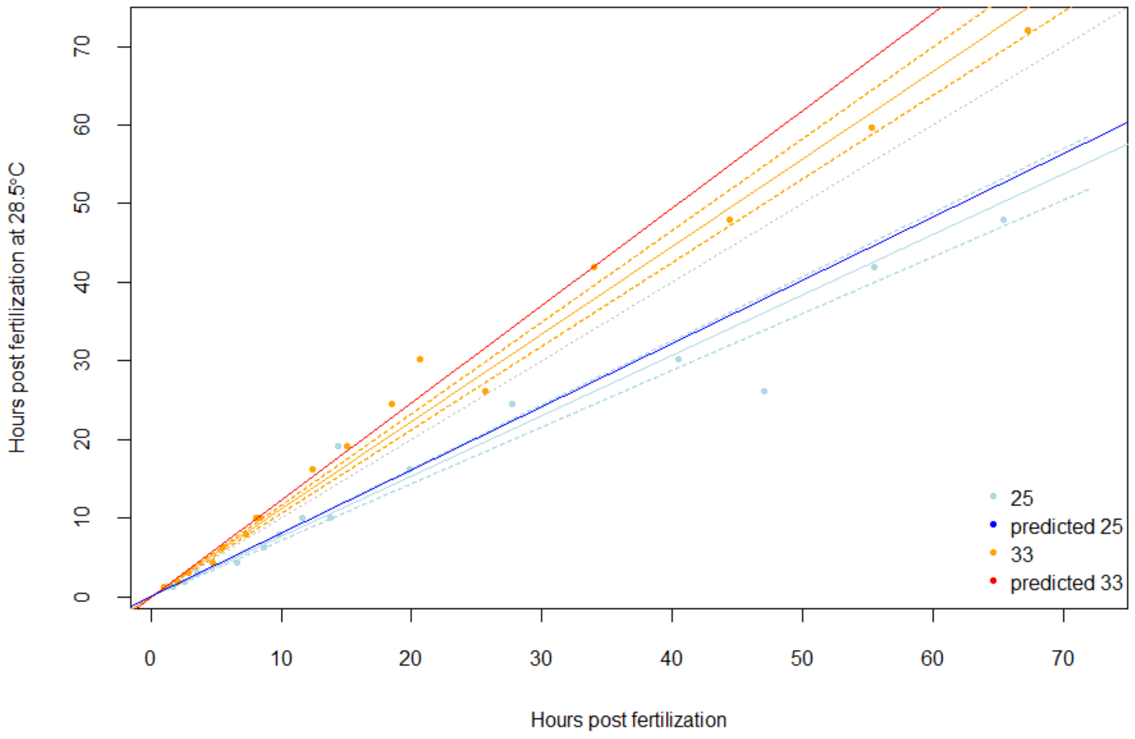


***Fig. S5:*** *Relationship between the hours post fertilization (hpf) at different temperatures and the corresponding normalized hpf at 28.5 °C, comparing observed data points with predicted values based on fitted regression lines (adapted from Kimmel et al. (1995)). Raw data is listed in Table S5.*

***Table S5:*** *Raw data of developmental rates derived from Kimmel et al. (1995) by using Web Plot Digitizer (Raharzi, 2024). Shown are the hpf at 28.5 °C, which is the standard incubation temperature, and the development of the eleutheroembryos in hpf when incubated at higher or lower temperatures.*

| hpf at 25 °C | hpf at 28.5 °C | hpf at 33 ° C |
|---|---|---|
| 1.7 | 1.3 | 1 |
| 2.6 | 2 | 2.1 |
| 3.6 | 3 | 2.9 |
| 6.6 | 4.3 | 4.7 |
| 8.7 | 6.3 | 5.5 |
| 9.8 | 8 | 7.3 |
| 11.6 | 10 | 8.1 |
| 13.8 | 10.1 | 8.4 |
| 14.4 | 16.3 | 12.4 |
| 19.9 | 19.2 | 15.0 |
| 27.7 | 24.5 | 18.5 |
| 40.5 | 26.1 | 20.6 |
| 47.0 | 30.2 | 25.6 |
| 55.5 | 41.9 | 34.0 |
| 65.4 | 47.9 | 44.4 |
| 76.7 | 59.7 | 55.3 |
| 87.9 | 72.1 | 67.3 |
